# A low-cost smart system for electrophoresis-based nucleic acids detection at the visible spectrum

**DOI:** 10.1101/2020.04.26.062604

**Authors:** Eduardo Nogueira Cunha, Maria Fernanda Bezerra de Souza, Daniel Carlos Ferreira Lanza, João Paulo Matos Santos Lima

**Affiliations:** Programa de Pós-graduação em Bioinformática (PPg-Bioinfo), Instituto Metrópole Digital (IMD), Universidade Federal do Rio Grande do Norte (UFRN); Programa de Pós-graduação em Bioquímica, Centro de Biociências, UFRN; Laboratório de Biologia Molecular Aplicada (LAPLIC), Departamento de Bioquímica, Centro de Biociências, UFRN; Laboratório de Sistemas Metabólicos e Bioinformática (LASIS), Departamento de Bioquímica, Centro de Biociências, UFRN; Instituto de Medicina Tropical do Rio Grande do Norte (IMT-RN), UFRN; Bioinformatics Multidisciplinary Environment (BioME), IMD, UFRN

**Keywords:** electrophoresis, low-cost, nucleic acid detection, IoT, LEDs

## Abstract

Nucleic acid detection by electrophoresis is still a quick and accessible technique for many diagnosis methods, primarily at research laboratories or at the point of care units. Standard protocols detect DNA/RNA molecules through specific bound chemical dyes using a UV-transilluminator or UV-photo documentation system. However, the acquisition costs and availability of these devices, mainly the ones with photography and internet connection capabilities, can be prohibitive, especially in developing countries public health units. Also, ultraviolet radiation is a common additional risk factor to professionals that use electrophoresis-based nucleic acid detection. With that in mind, this work describes the development of a low-cost DNA/RNA detection smart system capable of obtaining qualitative and semi-quantitative data from gel analysis. The proposed device explores the visible light absorption range of commonly used DNA/RNA dyes using readily available parts, and simple manufacturing processes, such as light-emitting diodes (LEDs) and 3D impression. By applying IoT techniques, our system covers a wide range of color spectrum in order to detect bands from various commercially used dyes, using Bluetooth communication and a smartphone for hardware control, image capturing, and sharing. The project also enables process scalability and has low manufacturing and maintenance costs. The use of LEDs at the visible spectrum can achieve very reproducible images, providing a high potential for rapid and point-of-care diagnostics as well as applications in several fields such as healthcare, agriculture, and aquaculture.

## INTRODUCTION

Despite constant growth and improvement of new technologies associated with molecular biology, PCR (Polymerase Chain Reaction) analysis still accounts for the largest share of the international market. The revenue expected for this area is estimated to be around 10.776 billion dollars by 2023, registering a CAGR (Compound Annual Growth Rate) of 6.2% from 2017 to 2023 (01)(02). Such growth of a consolidated technique shows opportunities for innovation of developing countries companies, including ones that will reduce local costs and the need for imported components, which will enable these technologies widely available.

Nucleic acid detection by gel electrophoresis is a ubiquitous laboratory routine procedure used in several fields, like genetics and molecular biology, biochemistry, and forensic science (03), and an essential step in extraction, cloning, and PCR workflows (04). In the last decades, its increasing application allied with more accurate molecular diagnosis techniques has provided useful information on the general condition of patients, as well as contributing to the diagnosis and prognosis of various diseases (05). Despite its simplicity, equipment and analysis costs are still high, especially in low resource settings (LRS) or for point-of-care (POC) applications.

The most used chemical dye for gel electrophoresis-based DNA/RNA detection is Ethidium Bromide (EtBr) (06). This dye bounds to DNA and, in the presence of ultraviolet (UV) light between wavelength 260 and 360 nm, fluoresces in orange-red range, in the length of 590 nm, being able to detect as little as 10 ng of DNA (07). However, the use of EtBr presents several disadvantages, among them, is the fact that it is highly mutagenic and carcinogenic, as well as its requirement for UV light exposure for detection. Non-toxic fluorescent dyes are good alternatives to EtBr (08) nucleic acid staining, such as Methylene Blue, which has no mutagenic potential and no UV light need, although it is significantly less sensitive (09) and Sybr Green, which exhibits about the same sensitivity as the EtBr and can be visualized in the blue or UV light. More recently, there is also a line of more efficient and safe staining-ready gels, such as GelRed and GelGreen. However, they are expensive alternatives, and most laboratories, especially in developing countries, still use EtBr, despite its disadvantages.

The use of EtBr also popularized UV transilluminators for the visualization of nucleic acid gel electrophoresis. This device generates light at UV wavelengths to excite the fluorophore present in the gel, allowing the visualization of the molecule to which it is attached. However, these wavelengths are not the same for all DNA/RNA specific dyes. Therefore, most equipment available on the market is not specific for all applications in research and clinical laboratories. Although it is a conceptually simple instrument, transilluminators have a high cost, about U$ 500.00 to 2,000.00, even without adding photodocumentary capabilities, which is usually only justified by its application. Moreover, camera-coupled transilluminators usually need to be connected to a computer and positioned on a transilluminator or have an integrated chamber for the acquisition of images. The associated costs, coupled with its low portability, often make it infeasible to use in places with limited structures such as LRS laboratories or POC health units in developing countries.

With the advancement of technology and its availability, it is possible to develop systems that previously had a very high cost at very competitive prices in the market. Also, the growth of “do it yourself” mentality, the increasing accessibility to top-end technologies, and the Internet of Things (IoT), all have broken paradigms of connectivity and development of electronic products (10). The most significant search today in all areas is for integrated, portable, low energy, automatic, and low-cost smart devices. In the laboratory environment, simple technologies combined with excellent engineering had allowed a reduction of up to 20 times in the values of some equipment. At present, easily accessible materials can allow the development of thermal cyclers for PCR (11), electrophoresis apparatus (12), and nucleic acid direct detection systems (13) (14) (15). With that in mind, the aim of the present work was the development of a low-cost DNA/RNA detection smart system capable of obtaining qualitative and semi-quantitative data from gel electrophoresis analysis from different available dyes. With easily accessible material and technologies, as light-emitting diodes (LEDs), miniaturized computing, and 3D impression, and also using a smartphone and custom software, it was possible to obtain a capable device to substitute a conventional commercial UV-transilluminator in several applications.

## MATERIALS AND METHODS

### System overview

The device development is based on IoT concepts and utilizes equipment and components that are readily available on the market. The priorities were to keep the low cost, but still have a design that is connected, easy to reproduce, and operate, and also capable of producing high-quality gel images. The project of the device comprises two parts, smartphone software and the system hardware per si (Fig 1), connected by Bluetooth protocols.

**Fig 01.**
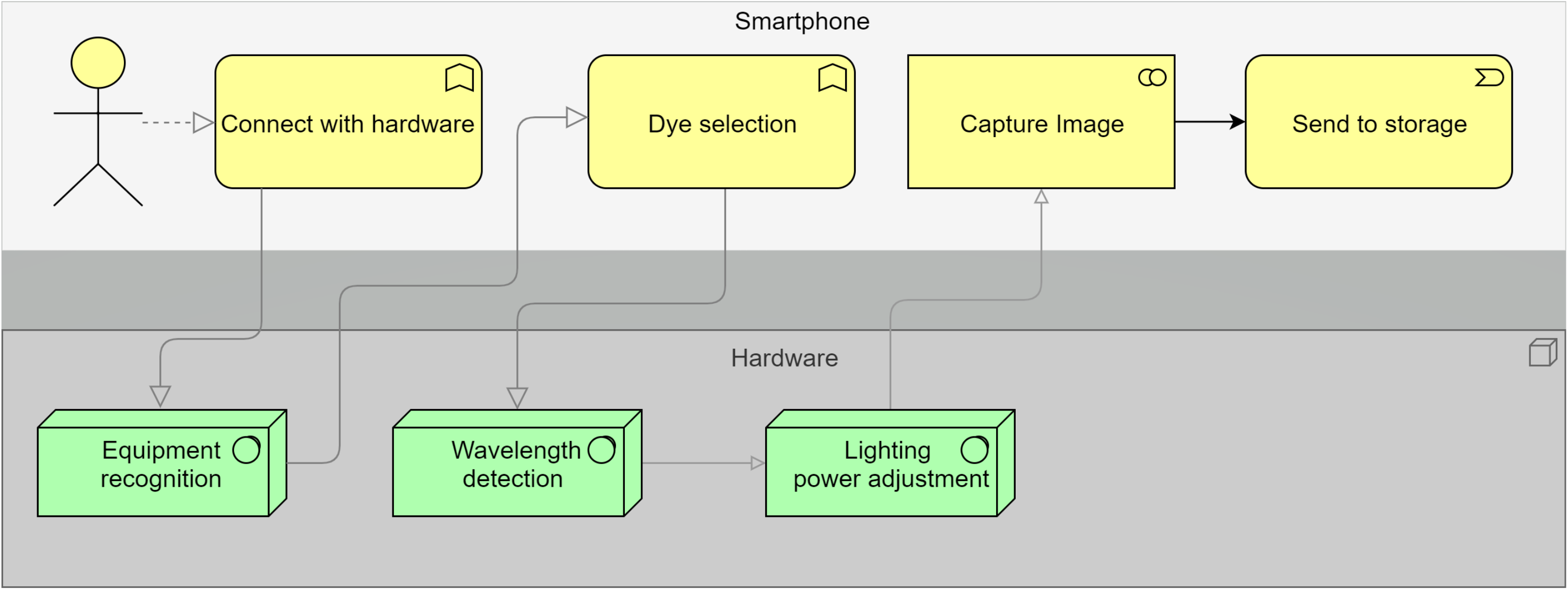
A schematic diagram demonstrating an overview of the model’s project. The diagram also describes the tasks executed by the smartphone software (yellow boxes) and by the hardware (green boxes), and their respective workflow.

*System hardware* - consists of a microcontroller platform with wifi and Bluetooth connections. It also includes an adaptable controller to various dye types, selecting the wavelength emitted by the lighting system for the type of dye used in the experiment, summarized in Table 1. Lighting intensity control is also implemented in hardware to optimize image capture, as shown in (16).

**Table 01.**
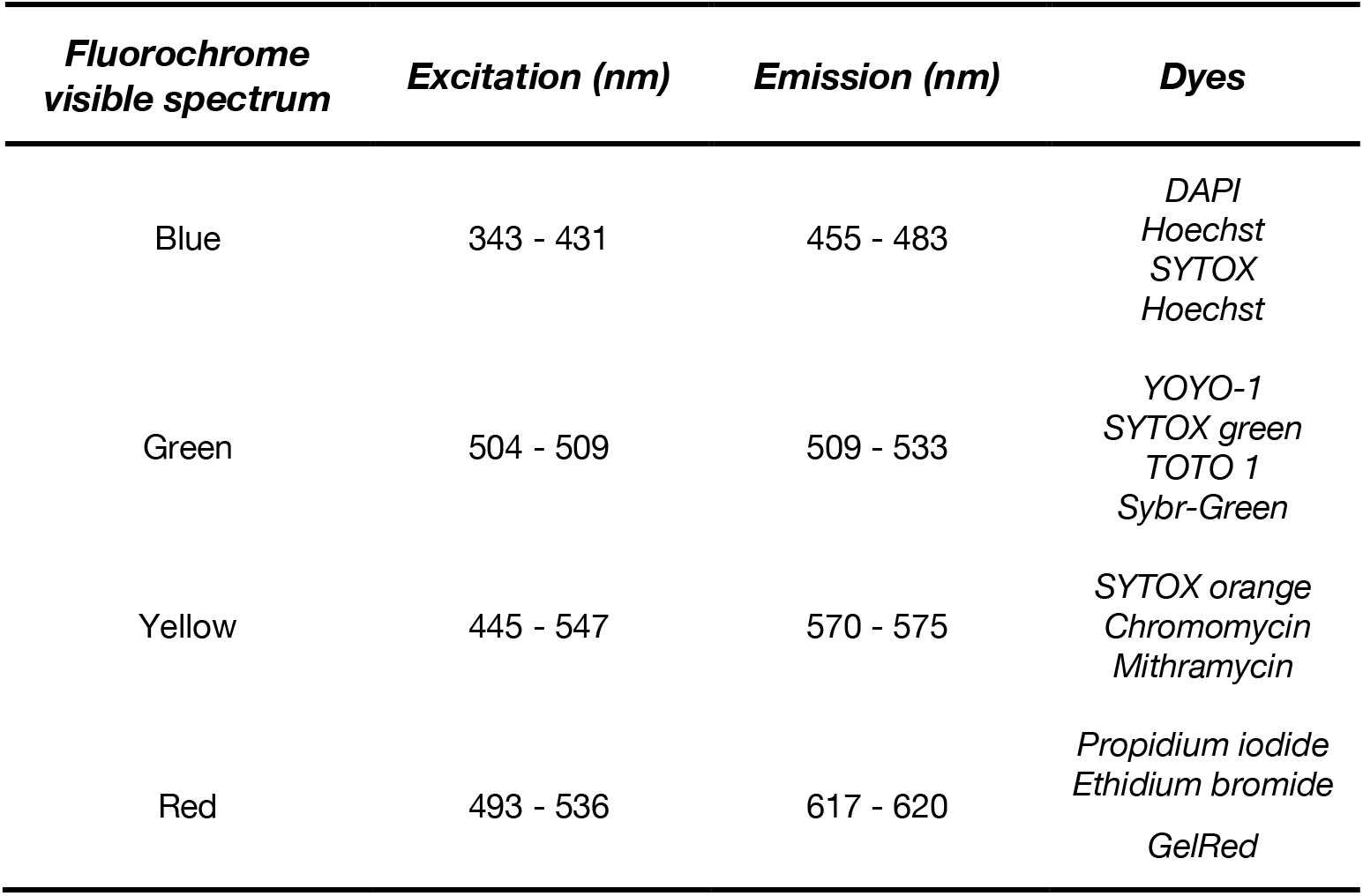
Excitation and emission wavelengths for commonly available commercial dyes. Reference guide to the detection of several dyes using LEDs under the visible light spectrum.

*Smartphone software* - developed for the Android platform, the software connects directly to the hardware via Bluetooth and performs image capture and fluorophore type selection stages, which can be automated using a wavelength scanning process. At this step, there is also an option to automatically or manually validate the captured image. Another function implemented in smartphone software along with hardware is image capture optimization (17)(18), to synchronize the camera frequency with the pulse cycle of the used LEDs. After the image validation process, the software sends data for storage or remote analysis.

### Hardware

#### Lighting system

For the excitation of the dyes, the system uses fourteen 3 W RGB LED chips, with a coupled heat sink, divided into three separate sets. Each set has unevenly powered red, blue, and green

LEDs. This division in the electric power does not happen in the same way (as will be described in the elaboration of the control circuit of the led) since the voltage of the red LED matrix is smaller. This uneven power division compensated the amount of lux emitted by the red LED and permitted an equal luminous power.

#### Electronic circuit

For the design of the LED’s control circuit, we used a pulse width modulation (PWM) system. To obtain control and optimize the lighting system according to the arrangement of the box, we used one PWM for each set of LEDs. The LED chip RGB consists of three parallel LEDs, mounted with a common anode, which need the same voltage (24 V) and current (600 mA) for each color matrix in each LED. The drive circuit has taken into account the switching speed for pulse modulation so that it is possible to obtain the full-color spectrum within the visible, and there is no excess energy dissipation by Joule effect (19), not compromising the durability of the card and components.

The microcontroller used was the ESP32 (Espressif) (20), operating at 0.8 mA current at the port, well below its maximum capacity of 12 mA. The system’s project also included touch screen buttons and signal operation LEDs mounted to an auxiliary circuitry. For controlling the LED cooling system, aluminum heat sinks were mounted directly on the LED board. The circuit contained correctly dimensioned components to avoid overload of the LED chips during use. Fig 02 describes the circuit project and shows the components already sized.

**Fig 02.**
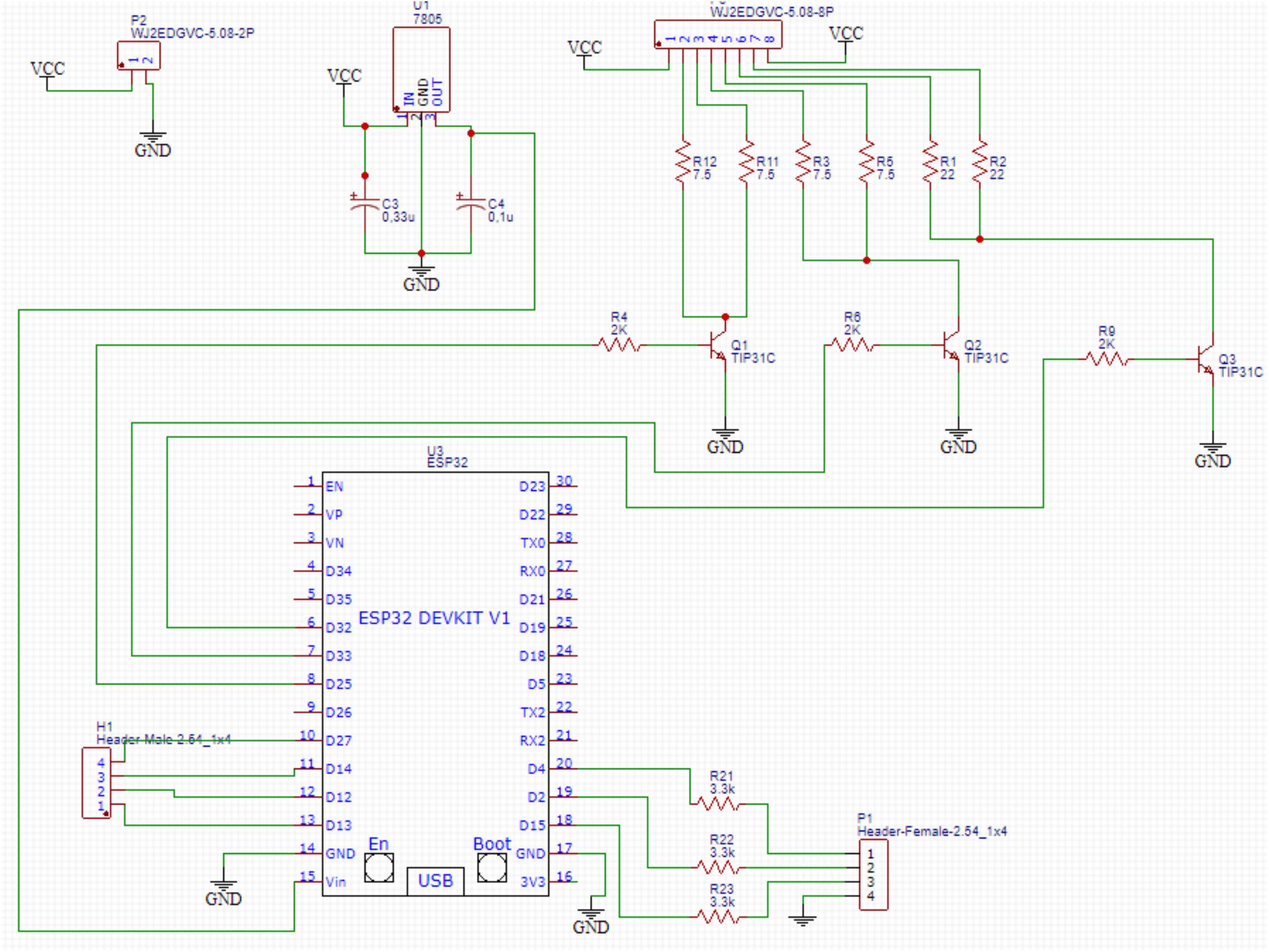
Diagram representing the system’s electric circuit, describing the component dimensioning and best arrangement, as well as the respective microcontroller inputs and outputs.

#### 3D printing

To maintain a low cost and achieve a straightforward reproduction, we opted to use 3D printing for the development of the platform’s on-site photographic documentation system. Fig 3 shows the final prototype. The calculation of the size and angles for the smartphone support depends on its camera’s specifications and must guarantee that the entire illuminated area is covered. In the tested prototype, we used the following measures:

**Fig 03.**
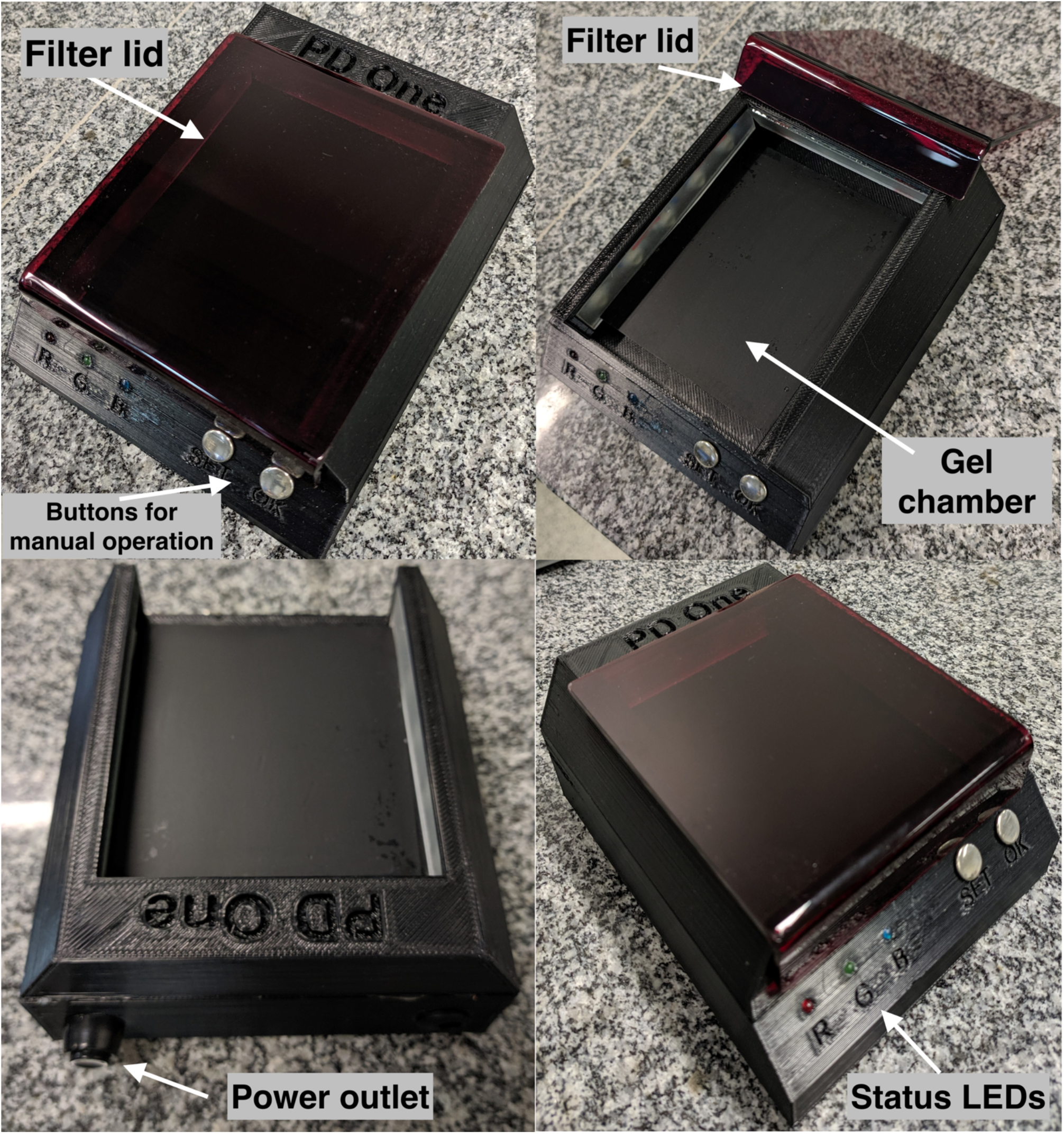
Final assembly of the system after 3D printing. White arrows indicate the filter lid, the status LEDs, the gel chamber, the power outlet, and the “SET” and “OK” buttons for manual operation of the system.

– Xiaomi MI 9 SE smartphone, the sensor Sony IMX586 Exmor RS, there is three cameras 48 Mp + 16 Mp + 12 Mp, resolution 8k x 6k pixel, sensor size 1/2 “ + 1/3 “ + 1/3.6 “ and aperture size F 1.75 + F 2.2 + F 2.2. For this experiment, we used only the 48 MP camera, but a minimum setting of 12 MP is sufficient for viewing the bands on the proposed device.
– The measures of the support.

– Height 120 mm, width 120 mm, and depth 145 mm.
– Minimum distance for image capture: 90 mm.

Supporting information includes simple calculations for the support’s measures, for standard cameras available. The constructed prototype can accommodate gels in any range according to the calculations presented. The inclusion of the SET and OK buttons allows the system’s operation as a simple transilluminator, without using a smartphone, though power and wavelength selections are software-managed only. Supplementary Fig 1 describes the CAD drawing model for 3D printing.

### Software

The system includes two operating software. The ESP32 microcontroller software, written in C programming language, has the function of controlling the pulse width modulation system of the LED chips, the communication with the smartphone via Bluetooth low energy (BLE), and the touch buttons. The Android smartphone application is responsible for connecting to the ESP32 microcontroller, as well as managing the type of dye used, its frequency (color) and intensity with the emitted light. This application, developed in an online block programming environment at Thunkable®, also controls the included photography and sharing functions.

### Sample preparations and gel electrophoresis

The template DNA samples used for electrophoresis were from the shrimp genomic DNA an 18S sequence of extremely conserved ribosomal DNA in decapod, the PCR product has a similar pattern in almost all decapod with an 848 bp amplifier, specific primer pair corresponds to nucleotide sequence 352–1200 of the 18S rRNA, using primers (143F 5’-TGC-CTT-ATC-AGCTNT-CGA-TTG-TAG-3’ and 145R 5’-TTC-AGN-TTT-GCA-ACC-ATA-CTT-CCC-3’, N represents G, A, T or C), described in (19). PCR’s reactions had a final volume of 75 µL containing: 0.25 µM from each primer, 2.5 mM MgCl_2_, 2.5 mM of each dNTP, 1 U GoTaq® DNA Polymerase (Promega) and 6 µL of the template DNA. PCR products were quantified using the QuantiFluor One dsDNA system. The concentration of the PCR final product was 38 ng/µL. Eight samples from seven serial dilutions of the PCR final product (38.00, 19.00, 9.50, 4.75, 2.38, 1.19, 0.59, 0.30 ng/uL) were prepared and later observed in three distinct 1% agarose gels, using SYBR Green®, GelRed®, and ethidium bromide as chemical dyes.

### System Calibration

In order to obtain the best viewing point for each of the dyes, we performed a calibration process using a tracking module implemented in the device’s software, which scans the entire region of the visible spectrum with an automatic increment of 10 nm (Fig 04). After that, we compared the results with the spectroscopy provided in each dye manual to ensure the use of its highest excitement regions. For the calibration process, ten independently prepared gels were used for each type of dye, with a more thorough scan for the dye with higher excitation within the UV region.

**Fig 04.**
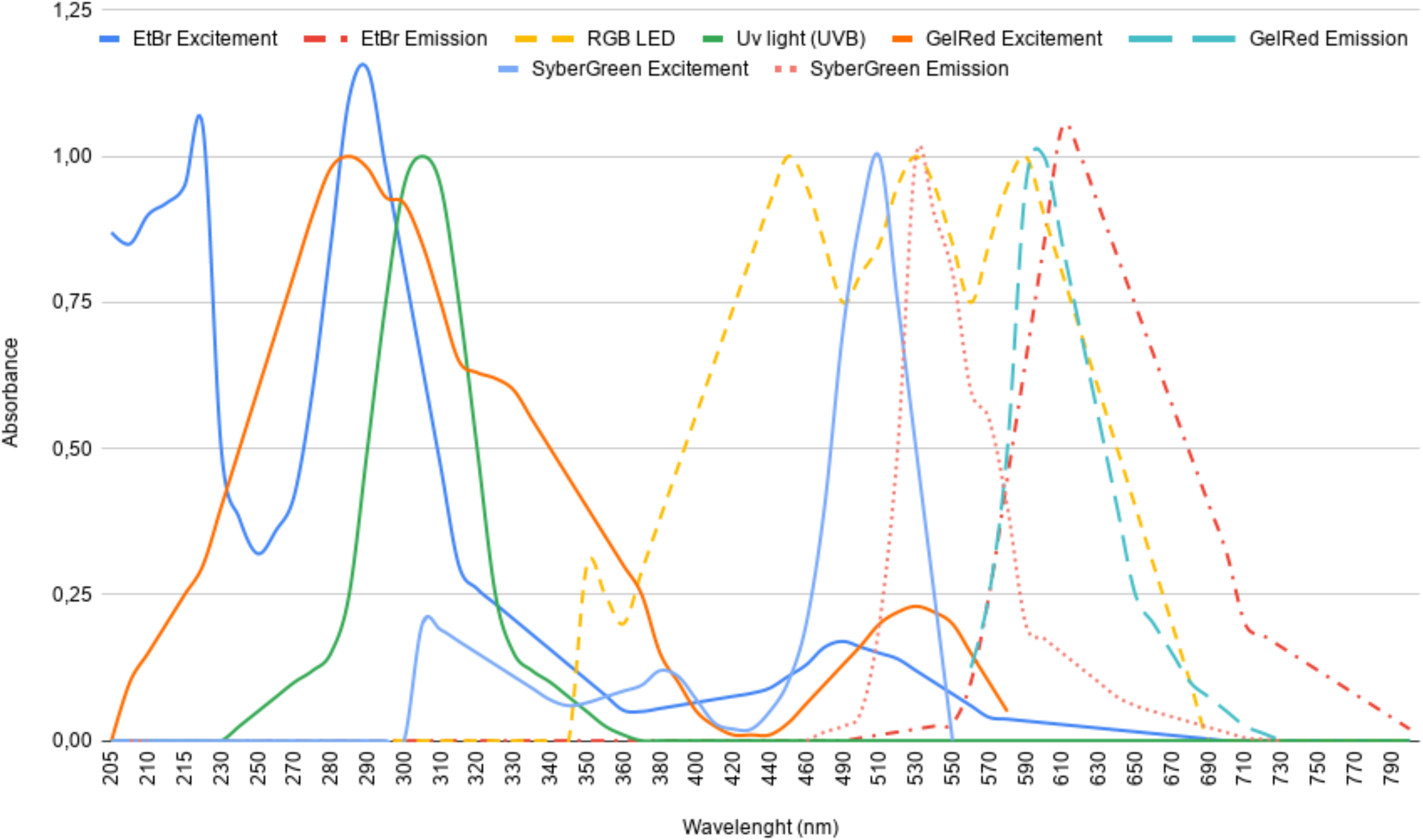
Plot representing the emission and excitation of the used dyes. Wavelengths are in the range of 205 - 790 nm (UV until visible). Full and dashed lines represent the emission and excitement wavelengths, respectively.

### Image analysis

During visualization within the system, all the gels were photographed with the coupled Xiaomi MI 9 SE smartphone (Android Version 10). For comparison reasons, the gels were also visualized and photographed using a 302 nm wavelength standard transilluminator. The resulting raw images (without software treatment) were analyzed and compared in ImageJ software (22).

## RESULTS AND DISCUSSION

### Assembly costs

We aimed to develop a complete device for DNA/RNA analysis using the electrophoresis technique, commonly used today in many research and health diagnostic laboratories. Table 2 describes the cost of each part. The average assembly cost was US$ 53.76, without including smartphone cost, which is pronouncedly cheaper than every commercially available system with similar functionalities. Rather than substituting a camera-coupled transilluminator entirely or for all its applications, this system is supposed to aid rapid and early electrophoresis-based diagnostics methods, especially in developing countries, either in POC units or at field or LRS conditions. Since there are different sizes of electrophoresis gels and their respective applications, the project presented here is easily modifiable to accommodate larger gels and can even be adapted to the construction of devices that use capillaries system for detection. Besides, by using a smartphone internet connection, results can be remotely sent to further analysis at reference health centers or laboratories. Regarding its low costs and uncomplicated manufacturer process, another potential application of this system is for science education at public schools, which usually have a low budget for scientific equipment (23).

**Table 02.** Parts and materials used in the manufacture and assembly of the system, and respective costs (not including the smartphone price).

| ***Part/Material*** | ***Quantity*** | ***Cost (US\$)\**** |
| --- | --- | --- |
| ESP32 microcontroller (Expressif) | 1 plate | 3.76 |
| LED RGB 3W (OS 8060C) | 14 LED chips | 8.00 |
| Acrylic (3mm thick red acrylic) | 80 cm<sup>2</sup> | 2.00 |
| 3D printer PLA filament (black color, 1,75 +-0,05 mm) | 300 g | 9.00 |
| Various Electronics components (resistor, transistors, capacitors, printed circuit board and connectors) | - | 10.00 |
| Various Miscellaneous Parts (masking tape and separators) | - | 1.00 |
| 3D printing (print time 13h) | 13h | 20 |
| **Total cost (without smartphone)** |  | **53.76** |

\*Average costs of the required materials. These costs refer to a system able to accommodate gels sizes from 80 cm<sup>2</sup>. The material acquisition occurred between the dates 08/01/2019 and 11/30/2019.

### Software management and dye detection

Using RGB LEDs, we were able to get all colors within the visible spectrum by combining the primary colors, red, green, and blue. Therefore, we got a piece of equipment that can excite any commercially available dye, since it has a known visible-light emission wavelength range. As can be evidenced in Fig 4, most DNA/RNA specific dyes have a low but sufficient excitement within the visible spectrum. For EtBr, a commonly used dye in molecular analysis laboratories, due to its low cost, we achieved excitation, emission, and detection without the use of UV light. Although EtBr has a higher excitement within the UVB (280 - 315 nm) region, conventional UV lamps used in transilluminators have a considerable displacement, not taking full advantage of this peak region (24), which is still significantly higher than the excitement region within the visible spectrum (25). However, our system can excite EtBr molecules at its peak visible absorption, around 480 nm, by using high-luminance LEDs, which obtain a better excitement optimization at this range and partially compensate for this energy difference. The developed smartphone associated software has a screen for selecting the dye used for detection (Fig 5). Afterward, it automatically sets LED colors to maximize dye-specific excitement. Also, since the tracking module scans the entire visible region, the adjustment of the system is possible to ensure the use of dyes’ highest excitement regions. Therefore, calibration is essential for further optimization and also for the correct registration of wavelengths for each dye in the smartphone software.

**Fig 05.**
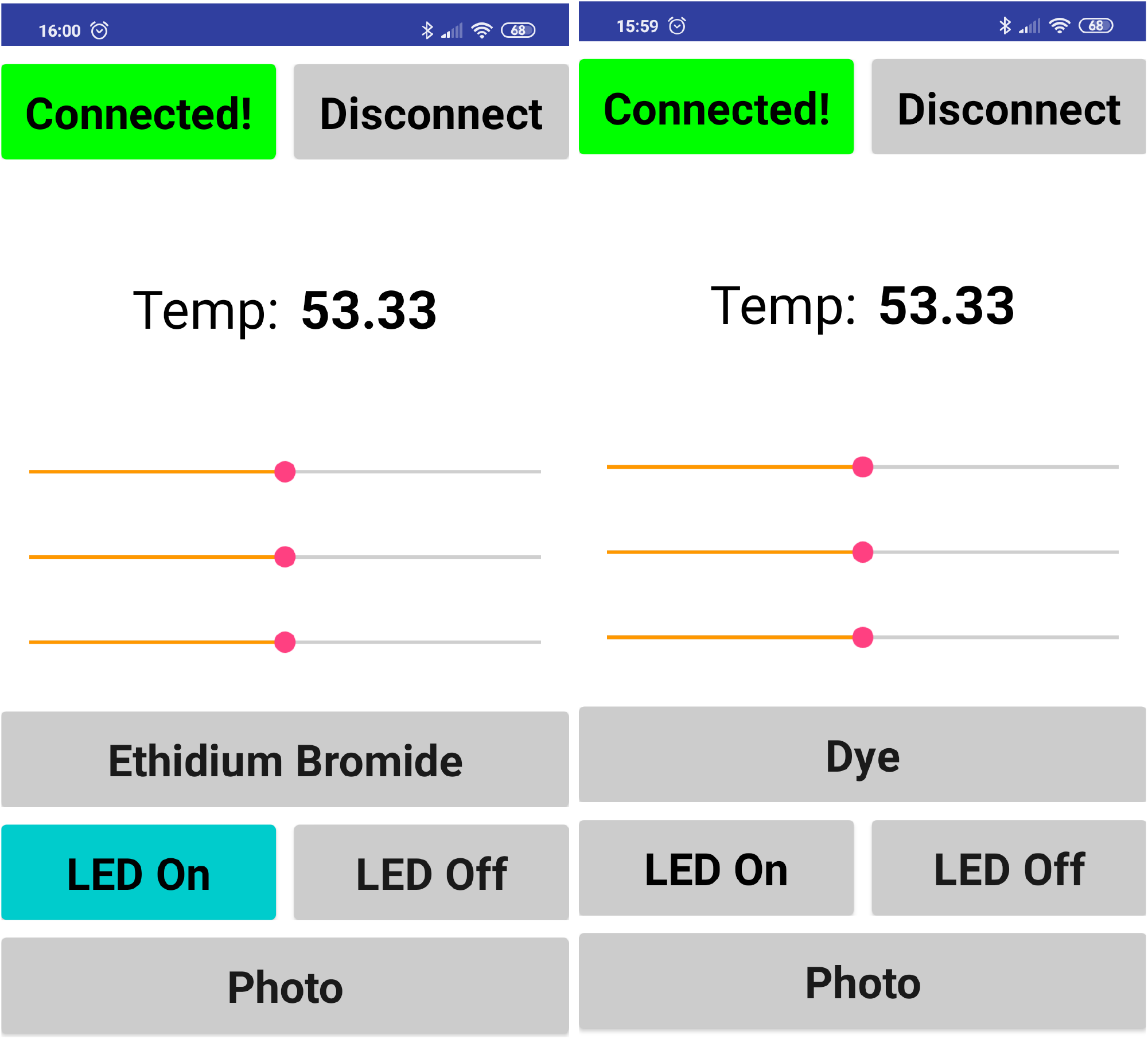
Screenshots of the smartphone software. In the software, it is possible to monitor the device temperature and to make the fine-tune adjustments for dye selection. Initial screen (right). Preset for using ethidium bromide as a dye (left).

### Sensitivity analysis

To evaluate the visualization potential of the developed system, we used the same amplified DNA sample dyed with three distinct and commonly used dyes (EtBr, gelRed, and SYBR Green), comparing the results with the visualization using a standard UV-transilluminator. By using smartphone software communication (Fig 5), we selected the highest point of excitation from each dye within the spectrum for the RGB LED, as shown in Fig 4.

As can be seen in Fig 6A and Fig 6B, the visualization of EtBr stained bands is more definite and precise in the photos taken on the UV-transilluminator. As discussed before, this is due to the high adsorption potential of this dye within the UV region (Fig 7). By analyzing the plot of the grayscale difference of bands’ densitometry (Fig 7) from the proposed device, we perceived that it is not possible to view the samples at concentrations below 3.0 ng/uL. We observed similar results for the gelRed staining (Fig 6 C, D)(Fig 7). However, despite the difficulty of visualization of the more diluted EtBr and gelRed-dyed bands (1.52, and 0.76 ng/uL) in photos taken by using the proposed device, it does not invalidate its use for these dyes. Low concentrations are usually not recommended for semi-quantitative analysis (26). Additionally, optimization of protocols and sample concentrations can compensate for this potential weakness, with the advantage of using a device that is safer than conventional UV-transilluminators, since it does not emit UV radiation.

**Fig 06.**
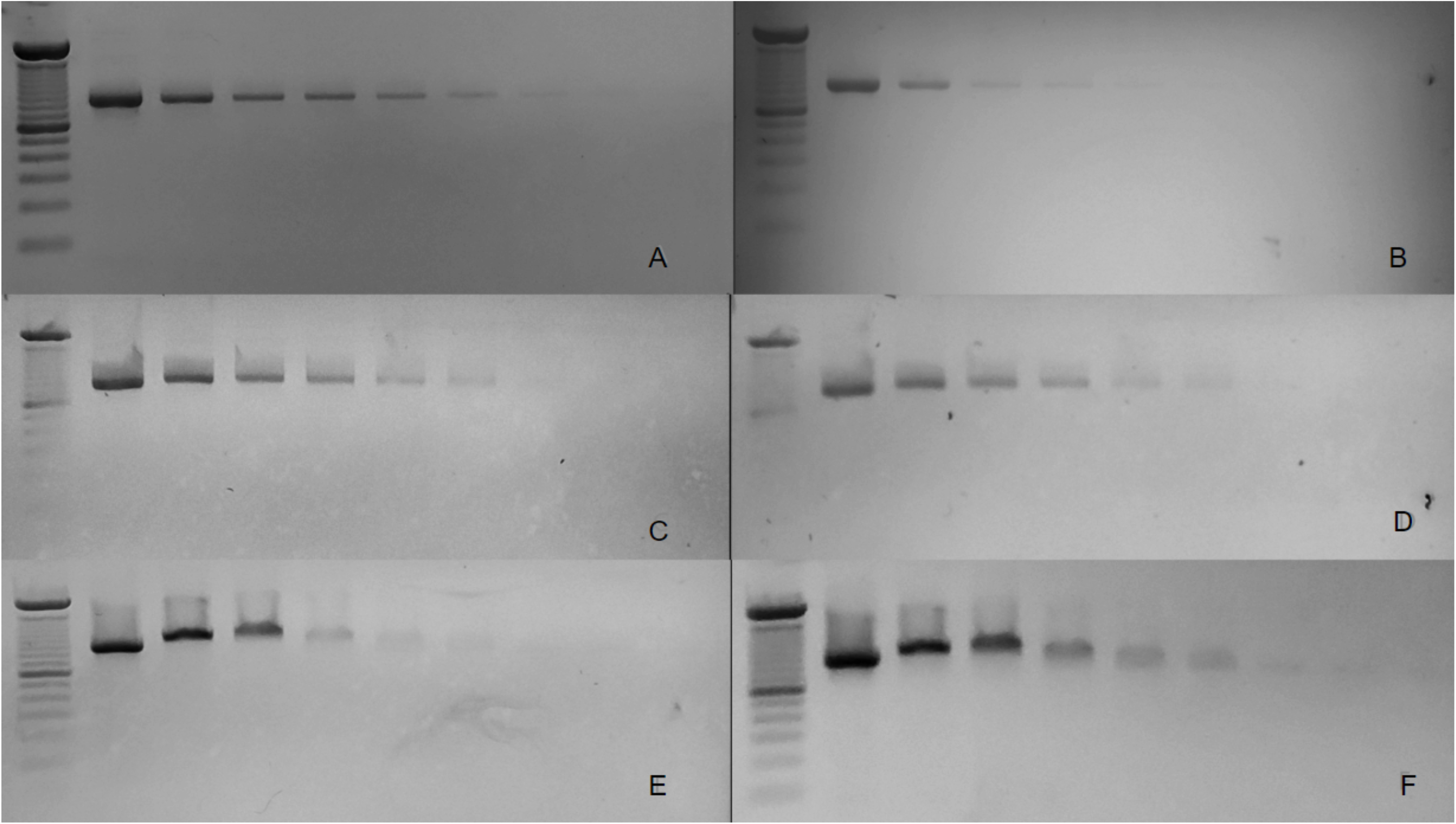
Agarose gel electrophoresis images from three different dyes. Ethidium bromide (A, B); Gel-red (C, D); and SYBR Green (E, F). Photos obtained from the use of a conventional UV-transilluminator are on the left panel (A, C, and E). Images captured from the proposed device are on the right panel (B, D, and F). All photos underwent the same image treatment in the ImageJ software (22) so that there was no advantage for comparative analysis. Samples in the gel lanes are serial dilutions of the PCR final product (see Material and Methods).

**Fig 07.**
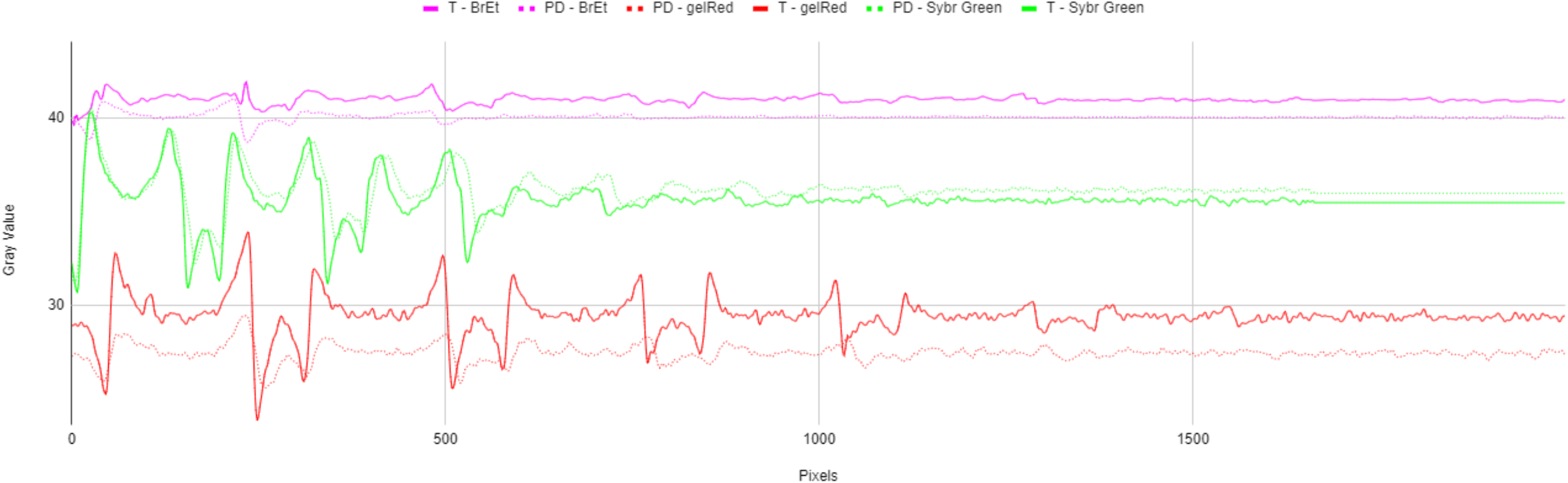
Gel images analysis. Analysis of intensity quantification (by pixel density) of the agaroses gel images obtained with a conventional transilluminator (full lines) and our proposed system (dashed lines), with the three used dyes: Ethidium bromide (purple lines); Gel-Red (red lines); Sybr green (green lines).

Nevertheless, the device surpasses conventional UV-transilluminators for the use of SYBR green-based protocols. SYBR-stained DNA bands were more brilliant and pronounced in photos taken under the illumination of visible LEDs (Fig 6E, 6F) (Fig 7). The system takes advantage of the high absorption potential of this dye within the visible region. Under this chemical staining, it is difficult to evidence samples dilutions below 3.0 ng/uL under the transilluminator radiation, the exact opposite from the results of the two other dyes. Although it also has its toxic effects (27), SYBR-green has surged as a safer alternative for EtBr nucleic acid staining (28), with the plus of having a more uncomplicated decontamination process (29).

### Functionalities and features of the system

Table 3 summarizes the main features and specifications of the proposed system compared to conventional commercial transilluminators. Our system is a low-cost example of a new generation of transilluminators that aims to apply the concepts of IoT in laboratory equipment. The insertion of the technology has the clear objective of increasing security both in not using UV radiation but also avoiding the user’s contact with the transilluminator, by the use of a smartphone as a control for dye adjustment and image capture and processing.

**Table 03.** Features and specifications comparison between a conventional commercial transilluminator and the proposed device.

| ***Part/Material*** | ***Commercial***<br>***Transilluminator*** | ***New***<br>***Transilluminator*** |
| --- | --- | --- |
| UV radiation | x |  |
| Adjust wavelength according to dye |  | x |
| Wireless operation |  | x |
| Estimated lamp life | 1000 h | 50.000 h |
| Energy consumption | 1,5 KW | 0,15 KW |
| Weight | 7 Kg | 0,4 Kg |
| Cost | U\$ 500-2000 | U\$ 53,76 |

## CONCLUSION

The low-cost smart detection system here presented can substitute a conventional UV-transilluminator in several electrophoresis-based applications, allowing the detection of nucleic acid bands staining with three commercial dyes. Besides the low-cost, other advantages of our system are the maximization of dye-specific excitement under the visible light spectrum, portability, and connectivity. The performed analysis also demonstrated that, in some cases, the device could achieve better results than a standard UV-transilluminator for the detection of SYBR green-dyed DNA bands. These features, allied to its easily accomplishable construction, confirm the potential use of this device in low resource settings (LRS) or for point-of-care (POC) and educational applications.

## SUPPORTING INFORMATION

Document file describing the calculations for the support’s measures, for standard cameras available (pdf).

## ACKNOWLEDGEMENTS

The authors would like to thank the *Núcleo de Pesquisa e Inovação em Tecnologia da Informação* (nPITI) and the Digital Metropolis Institute (IMD) from the Federal University of Rio Grande do Norte (UFRN) for the support in the realization of this work.

## AUTHOR CONTRIBUTIONS

EN Cunha, DCF Lanza, and JPMS Lima conceptualized the work. EN Cunha designed the electric circuit, constructed the detection system, and wrote the smartphone software. DCF Lanza and MFB Souza performed the molecular biology experiments. EN Cunha, DCF Lanza, and JPMS Lima wrote the paper.

