## Supporting information for "A low-cost smart system for electrophoresis-based nucleic acids detection at the visible spectrum"

### Supplementary Information

#### Calculations for equipment sizing

For the prototype design, a table was made based on tests of absorption and reflection of light at the illuminated area. All dimensions were predicted according to the area to be occupied by the gel. In Fig 01, Fig 02, and Fig 03 (from the main manuscript), we can see the prototype sized for a 70 x 80 mm gel. All measurements used in the formulas are in millimeters (mm).

- $Number\ of\ Leds\ =\ abs\left[\frac{With\ +Deph\ x\ 2}{20}\right]$
- $Led\ light\ angle\ of\ incidence\ = tg\left(\frac{gel\ width}{4}\right)^{-1}$
- $Minimum\ distance\ for\ image\ capture\ = \frac{tg\frac{\theta}{2}}{area}$
- $Equipment\ depth\ = gels\ depth\ + 45$
- $Equipment\ width\ = gels\ width\ + 40$
- $Equipment\ height\ = 56$

where:
$\theta = camera\ angles$;
$Area = gel\ area.$

#### Circuit Design

The circuit design has the aim to obtain the lowest possible current at the source. For this, we make a series of parallel arrangements with the LEDs (Fig 04, main manuscript), respecting the maximum current allowed by each LED. Another purpose of the circuit is to compensate for the difference between the luminous power versus the electrical power of the LEDs to obtain the best balance for capturing images.
